# Interrogating the Precancerous Evolution of Pathway Dysfunction in Lung Squamous Cell Carcinoma Using XTABLE

**DOI:** 10.1101/2022.05.06.490640

**Authors:** Matthew Roberts, Julia Ogden, A S Md Mukarram Hossain, Alastair Kerr, Caroline Dive, Jennifer E Beane, Carlos Lopez-Garcia

## Abstract

Lung squamous cell carcinoma (LUSC) is a type of lung cancer with a dismal prognosis that lacks adequate therapies and actionable targets. This disease is characterized by a sequence of low and high-grade preinvasive stages with increasing probability of malignant progression. Increasing our knowledge about the biology of these premalignant lesions (PMLs) is necessary to design new methods of early detection and prevention, and to identify the molecular processes that are key for malignant progression. To facilitate this research, we have designed XTABLE, an open-source application that integrates the most extensive transcriptomic databases of PMLs published so far. With this tool, users can stratify samples using multiple parameters and interrogate PML biology in multiple manners, such as two and multiple group comparisons, interrogation of genes of interests and transcriptional signatures. Using XTABLE, we have carried out a comparative study of the potential role of chromosomal instability scores as biomarkers of PML progression and mapped the onset of the most relevant LUSC pathways to the sequence of LUSC developmental stages. XTABLE will critically facilitate new research for the identification of early detection biomarkers and acquire a better understanding of the LUSC precancerous stages.

## INTRODUCTION

Lung squamous cell carcinoma (LUSC) is a type of non-small cell lung cancer that accounts for 40% of all lung cancer cases (1). Despite being the second most frequent type of lung cancer (2), our knowledge regarding the biology of this disease as well as the therapeutic modalities to treat it remain far behind the most frequent type of lung cancer, lung adenocarcinoma (LUAD) (3, 4).

LUAD genetics is dominated by mutations (that are often druggable) that activate the RTK/RAS pathway, including EGFR and KRAS mutations (5, 6). However, the genetic landscape of LUSC is more complex, with multiple pathways altered in subsets of patients and a lack of actionable mutations (7-9), precluding the development of new therapies. Hence, the only pharmacological therapies available to treat LUSC patients are immune checkpoint inhibitor monotherapy or in combination with chemotherapy (10-12). Furthermore, the National Lung Cancer Matrix Trial and The Lung Master Protocol, the largest personalized medicine trials in lung cancer, have not shown clear therapeutic benefits with targeted agents in LUSC (13-15). LUSC is also a more aggressive disease than LUAD, with a 5-year overall survival (OS) of 5% for patients diagnosed with distant metastasis (9% for LUAD). However, patients diagnosed with localized disease are eligible for curative surgery and the 5-year OS is 50% (16). Therefore, early detection is currently the most valuable tool to prevent deaths by LUSC as evidenced by several ongoing programs of lung cancer early detection (17-19). These initiatives make use of low dose CT-scans in high-risk populations, but in spite of the frequent detection of localized lung cancer eligible for resection with curative intent, 40% of early diagnosed patients die within 5 years.

LUSC progresses through a series of premalignant stages characterized by alterations of the normal bronchial epithelium (Figure 1) (20-23). These endobronchial premalignant lesions (PMLs) are classified as low-grade (squamous metaplasia, mild and moderate dysplasia) and high-grade (severe dysplasia, and carcinomas in-situ) (Figure 1). However, not all PMLs progress to LUSC. Although obvious differences exist between the multiple studies published on the topic, high-grade PMLs have a higher risk of progression than low grade, and high levels of chromosomal instability (CIN) are also predictive of progressive PMLs (22, 24-26). This potential role of CIN as biomarker of PMLs progression has been observed in low and high grade PMLs (24, 25). These reports showed that high levels of copy number variations was the best predictor of progressive PMLs (24, 25) and that immune response is the most likely cause of regression as these lesions contained higher levels of immune infiltration (23, 27). These results are supported by transcriptomic analysis of PMLs that indicate immune evasion in the transition to invasiveness (28).

**Figure 1:**
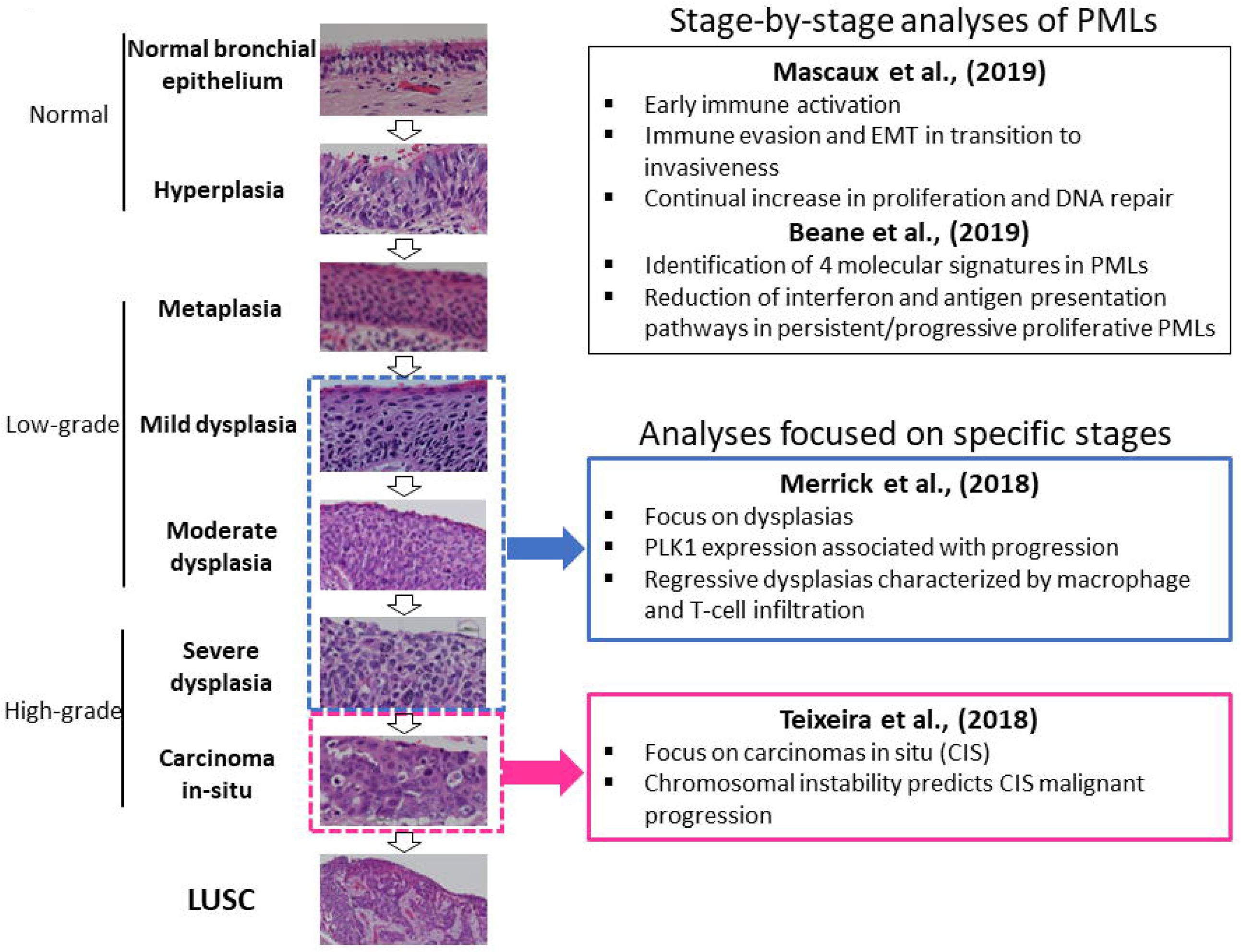
Developmental stages of LUSC PMLs with representative histological images for each stage (haematoxilyn-eosin) and a summary of the four studies included in XTABLE. PMLs are typically classified as normal epithelium (including hyperplasia), low-grade and high-grade. Two studies (Mascaux et al., 2019 and Beane et al., 2019) carried out gene expression analysis of multiple developmental stages, whereas Merrick et al., (2018) and Teixeira et al., (2019) focused on dysplasias (blue boxes) and CIS (pink boxes) respectively. The most relevant findings of each article are summarized in the figure.

The detection of PMLs cannot be carried out by routine patient imaging techniques such as CT or PET scans as the morphological change that they cause in the airway does not result in radiological contrast. Alternatively, ablation of high-grade PMLs detected by autofluorescence bronchoscopy using minimally invasive endoscopic procedures in high-risk populations is an innovative and interesting strategy to prevent LUSC (29). Nonetheless autofluorescence bronchoscopy is an expensive and complex technique of limited use in large screening programs. Therefore, simpler, more cost effective and scalable methods of high-grade PML detection are needed to prevent deaths by LUSC. Improving the detection of PMLs requires a better understanding of their biology and the validation of adequate biomarkers of progressive lesions that can be translated into new technologies for large screening initiatives. Cell surface proteins, metabolites, nasal-based biomarkers, blood and sputum/bronchoalveolar lavage biomarkers are examples of biomarkers that can be used to improve and/or complement current diagnostic techniques (such as CT and PET scans) or develop new ones.

Recently, scientific interest in the biology of preinvasive LUSC stages has motivated the publication of several articles characterising PML transcriptomes from various perspectives (Figure 1). Two reports published by Beane et al., (2019) (27) and Mascaux et al., (2019) (28) showed stage-by-stage gene expression analyses of PMLs. These studies provided the most detailed transcriptomic characterization of all PML stages so far and identified changes in the immune microenvironment associated with invasive transformation. Additionally, longitudinal studies by Beane et al., (2019) (27), Merrick et al., (2018) (30) and Teixeira et al., (2020) (25) contained samples with known progression potential. Beane et al., (2019) identified molecular subtypes with specific biological traits whereas articles published by Merrick et al., (2018) (30) and Teixeira et al., (2020) (25) focused on gene-expression and genomic analyses of specific preinvasive stages (dysplasias and carcinomas in-situ, respectively) with the objective of identifying predictors of PML progression and the biomolecular processes involved (Figure 1). These studies provide a valuable source of gene expression data to identify candidate biomarkers for the detection of high-risk PMLs and/or early stage LUSC as well as to investigate the biology of premalignant LUSC progression.

Simple and straightforward access to transcriptomic databases is a key requirement for the open science philosophy. Applications that integrate multiple databases allow cross comparisons between independent studies, strengthen the robustness of results obtained and allow the selection of high-confidence data. In this report, we provide an overall description of XTABLE (E**x**ploring **T**r**a**nscriptomes of **B**ronchial **L**esions) and provide examples of its functions. XTABLE is a new open-source application that will enable scientists to interrogate four LUSC PML transcriptomic datasets in a versatile manner that can be adapted to the needs of each researcher. Specifically, LUSC prevention and diagnosis are the main areas that can benefit the most from XTABLE, but its versatility and multiple functions lend themselves to the exploration of a variety of research questions. Without XTABLE, researchers would have to put together all the packages, data processing steps themselves for each analysis they wished to run. In this report, we provide an overall explanation of all the functions of XTABLE as well as a detailed description of the most important analysis modules, including two group comparisons, gene-of-interest analyses and interrogation of transcriptional signatures. Additionally, we explored the use of CIN-related signatures as a biomarker of progressive PMLs and mapped the onset of the most important LUSC pathways to its developmental stages.

## RESULTS

### Description of the studies included in XTABLE

Four datasets originating from four independent PML transcriptomic studies have been included in the XTABLE application (Figure 1, Table 1). Two datasets, GSE33479 (28) and GSE109743 (27), provide gene expression data of the developmental LUSC stages (with and without progression status information, respectively), whereas the remaining studies, GSE114489 (30) and GSE108124 (25) focus on analysing specific PML stages (dysplasias and carcinomas in-situ respectively) that have been followed up to establish their progressive, persistent or regressive potential.

**Table 1.** Description of the four cohorts included in XTABLE. (*) This cohort includes neither CIS nor Invasive carcinomas. Bx: biopsies; Br:brushings

| GEO accession | PMID | Stages | Progression status known | Number of samples | Sample type | Transcriptome |
| --- | --- | --- | --- | --- | --- | --- |
| GSE33479 | 31243362 | Multiple | No | 122 | Whole biopsies | Microarray |
| GSE109743 | 31015447 | Multiple* | Yes | 448<br>Discovery cohort (197 Bx, 91 Br)<br>Validation cohort (111Bx, 49 Br) | Whole biopsies and brushings | RNAseq |
| GSE114489 | 29997230 | Dysplasias Normal | Yes | 63 | Whole biopsies | Microarray |
| GSE108124 | 30664780 | CIS | Yes | 33 | Microdissected | Microarray |

Dataset GSE33479 (Mascaux et al., 2019) comprises expression microarray analyses of 122 endobronchial biopsies of unknown progression status. Samples were obtained following autofluorescence bronchoscopy and as no enrichment or microdissection of the biopsy epithelium was carried out, variable levels of stromal component are present in samples. Although this results in dilution of epithelial signals, it has the advantage of providing information about the microenvironment as well as insights into the infiltrated immune cells. Dataset GSE109743 (Beane et al., 2019) consists of a transcriptomic analysis of whole PML biopsies with no purification of the epithelial compartment as well as bronchial brushings obtained from adjacent normal regions of the bronchial mucosa. Unlike GSE33479, GSE109743 used RNAseq and more importantly, includes the progression status for some lesions established by serial biopsies. Samples were classified as ‘normal-stable’ when they changed between normal, hyperplasia and metaplasia, ‘regressive’ when they regress from dysplasia to a less severe dysplastic grade, or from dysplasia to normal/hyperplasia/metaplasia. Remaining samples were considered persistent/progressive. One advantage of this dataset is the large number of samples divided into a discovery and a validation cohort (Figure 1, Table 1). However, the representation of each PML stage is not homogeneous and carcinomas in-situ (CIS) samples are not included.

Datasets GSE114489 (Merrick et al., 2018) (30) and GSE108124 (Teixeira et al., 2019) (25) constitute a different type of study. Both make use of sequential biopsies to classify lesions according to their progression potential, but they differ in the stages included, the classification of progression status and sample processing. In GSE114489, the authors collected 63 baseline bronchial biopsies (with corresponding follow-up biopsies) and classified samples in four groups: 23 persistent dysplasias (dysplastic lesions with the same or higher severity scores in follow-up biopsies), 15 regressive dysplasias (dysplasias progressing to lower severity scores), 9 progressive non-dysplasias (biopsies with normal or hyperplastic morphologies that progress to more severe morphologies) and 16 stable non-dysplasias (normal or hyperplastic pathology that remain stable in follow-up biopsies. Microarray analysis of gene expression was performed on whole biopsies. The study published by Teixeira et al., (2019) (25) (GSE108124) also makes use of follow up biopsies to classify the progression potential of PMLs, but unlike GSE114489, GSE108124 focuses on CIS and progression is defined as the transition to invasive carcinomas in follow up biopsies. Whole-exome and methylation data are also available for some of the RNA-profiled samples in this study. RNA for gene-expression microarray analysis was extracted from microdissected FFPE samples to enrich the epithelial component.

### XTABLE access, interface and functions

XTABLE download can be carried out from https://gitlab.com/cruk-mi/xtable following the instructions in the methods section and Supplemental Video 1. XTABLE has been designed using the shiny app interface (see methods section) and its functions have been divided into 11 interrelated tabs that contain a specific function to interrogate each dataset separately (Figure 2A).

**Figure 2.**
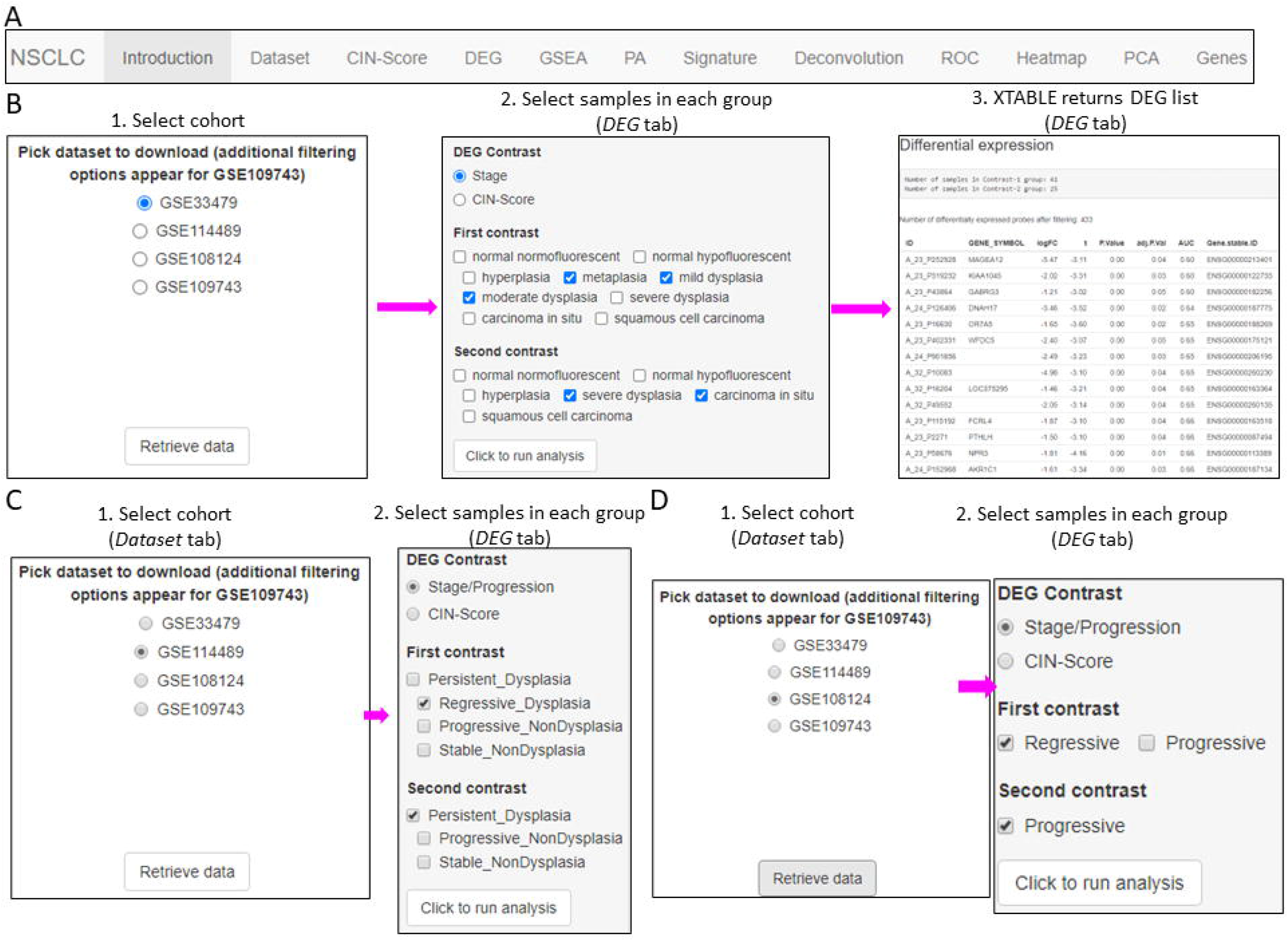
Overall organization of XTABLE functions and use of the *DEG* function. **A**. Organisation of all the functions in the XTABLE interface. The functions are interrelated and completing certain analyses requires the use of several functions. For instance, the *GSEA* and *PA* functions operate with gene lists obtained with the *DEG* function. **B**. Workflow to obtain differentially expressed genes between two groups using the *DEG* function. The example shows groups of samples arranged by developmental stage to compare low-grade and high-grade PMLs in the GSE33479 cohort. **C and D:** Workflow to obtain differentially expressed genes between two groups using the *DEG* function. The two groups have been arranged by progression status using in the GSE114489 and GSE108124 cohorts respectively.

#### Introduction

Detailed description of analyses performed in each tab, output formats, links to the 4 articles used in the application, bioinformatic packages used in the different functions and additional references.

#### *Dataset* tab

In this tab, the user selects the database (GSE114489, GSE108124, GSE109743 or GSE33479) for subsequent interrogation. Due to specific differences between the four studies, only one dataset can be interrogated at a time. Since dataset GSE109743 contains a discovery and a validation cohort, as well as biopsies and brushings, preselection of the cohort and type of sample must be carried out (Figure 2-figure supplement 1).

#### *CIN-score* tab

Chromosomal instability (CIN) has been identified as a good predictor of PML progression. Except for Teixeira et al., (2019), (25) the other three studies do not provide genomic analyses that provide an estimate of the level of chromosomal alterations in the samples. Several gene expression signatures that correlate with CIN (CIN-scores) have been described in the literature (25, 31). This *CIN-score* function returns a list of several CIN-scores (CIN70, CIN25 and CIN5, depending on the number of genes included in the signature) for all samples included in the study and a graph depicting CIN-scores in different sample types classified according to the sample types defined in the study. A line marking a selected CIN-score threshold can be added for visualization purpose (Figure 2-figure supplement 2).

#### *DEG* tab

Function to identify differentially expressed genes in comparisons of two groups of samples determined by the user. This function allows the selection of p-value and fold-change cut offs for the analysis.

#### *GSEA* and *PA*

In these two tabs, the user can carry out gene-set and pathway enrichment analyses using multiple tools (goseq, fgsea/MSigDB, enrichR, gage, kegga/pathview, ReactomePA, Progeny and Dorothea). This function operates with the list of differentially expressed genes obtained with the *DEG* function.

#### Signature

This function returns the gene-expression values of a user defined gene list. Lists can be manually entered or uploaded from a .csv file.

#### Deconvolution

Estimation of immune and stromal component in samples from gene-expression data using the ESTIMATE tool.

#### ROC

This function returns Receiver Operating Characteristic (ROC) curves in a user defined comparison of two groups of samples stratified by the expression of a gene or by CIN-scores.

#### Heatmap

Returns a gene-expression heatmap for genes selected by variance, user-defined gene signatures and differentially expressed genes in the *DEG* tab.

#### PCA

Principal Component Analysis of all the samples included in each study. Several sample characteristics can be highlighted in the PCA plot including CIN signatures, progression status and PML stage.

#### Gene

Statistical analysis of expression of individual genes selected by the user. Several options for sample groupings are available.

### Two-group differential expression analysis

The discovery of candidate biomarkers for the detection of PMLs at high-risk of malignant progression and the interrogation of PML biology depends greatly on the comparison of gene expression profiles between lesions with known progressive or regressive potential. The four databases included in XTABLE contain different types of information which influence how users can interrogate these databases. To facilitate the interrogation of the four transcriptomic databases in a manner that allows the versatile stratification of samples, we have designed a module in XTABLE named *DEG* (Figure 2B-D), that returns the differentially expressed genes in two user-defined groups of samples stratified by PML stage (GSE33479 and GSE109743), by known progression status (GSE109743, GSE114489 and GSE108124) or by CIN-score thresholds. In XTABLE, we have included three CIN-scores, named CIN70, CIN25 (31) and CIN5 (25). CIN70 and CIN25 have been reported in the literature, with the former containing the greatest number of genes for interrogation. CIN5 is derived from the signature used by Teixeira et al., (2019) (25) and is reported to show a good correlation with progression in CIS lesions.

Setting contrast groups by stage using *DEG* allows the comparison of two individual PML stages or the grouping of multiples stages into two groups. For instance, to compare the differential expression between low-grade and high-grade PMLs, we can define two contrast groups, one including metaplasia, low and moderate dysplasia (low-grade) and one including severe dysplasia and CIS (high-grade) using cohort GSE33479 (Figure 2B). After selection of the cohort and setting up the groups (Figure 2B), the application returns a downloadable list of differentially expressed genes with associated statistical information (Figure 2B), including AUC inferred by ROC analysis. ROC curves associated with each gene in a two-groups comparison, useful for biomarker discovery, can be downloaded using the *ROC* tab (Figure 2-figure supplement 3). Straightforward grouping of progressive and regressive samples can also be carried out by selecting the ‘Progression’ contrast in studies that provide this information (Figure 2C and D). To carry out two-group comparison by CIN signatures, the *CIN-score* tab provides a graphic visualization of the three CIN scores for all samples to guide the selection of a CIN-score threshold for sample stratification (Figure 3A). A .*csv* file containing the CIN-scores of all samples can be downloaded. The selected CIN threshold can be used in the *DEG* tab to stratify CIN-high and low samples and retrieve a list of differentially expressed genes (Figure 3A).

**Figure 3.**
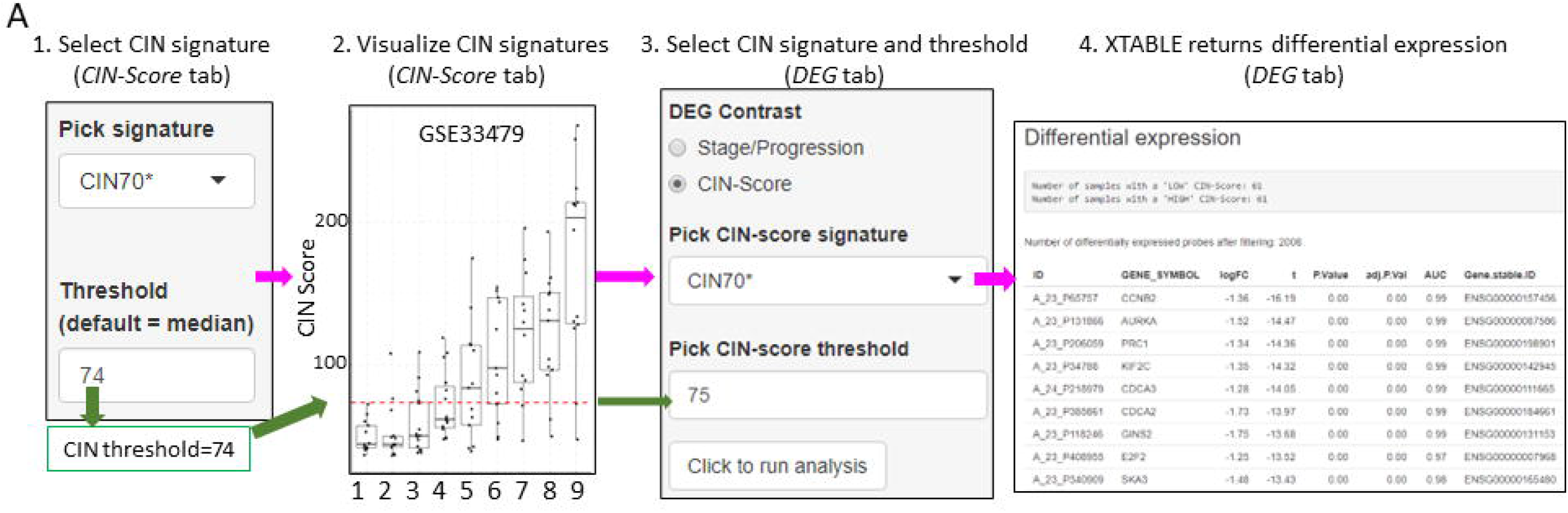
Differential expression analysis between two groups of samples classified according to a CIN-score threshold. The *CIN-score* function allows the graphic visualization of CIN-scores for all samples in a study. A CIN-score threshold selected by the user can be depicted on the graph (red dotted line). The CIN-score threshold selected by the user can be used in the *DEG* tab to define the two-group comparison. Stages 1 to 9 represent the 9 developmental stages of LUSC as described in Mascaux et al., (2019) (GSE33479). CIN70, CIN25 and CIN5 can be used in the *DEG* tab.

The list of differentially expressed genes obtained in the *DEG* tab, can be automatically used in the *GSEA* and *PA* tabs to carry out gene set enrichment and pathway analyses. The *GSEA* tab allows the user to select the gene set enrichment tool (goseq, fgsea/MSigDB, enrichR and gage), p-value cut-off and the gene sets to analyse (Figure 4A). For instance, using the goseq function enables us to select one of the three Gene Ontology domains (biological process, cellular compartment and molecular function) to consider for analysis (Figure 4A). Similarly, with the fgsea/MSigDB tool, the user can select the gene-set of interest (Figure 4B). The *PA* tab operates in a similar manner using four pathway analysis tools (kegga/pathview, ReactomePA, Progeny and Dorothea) (Figure 4-figure supplement 1).

**Figure 4.**
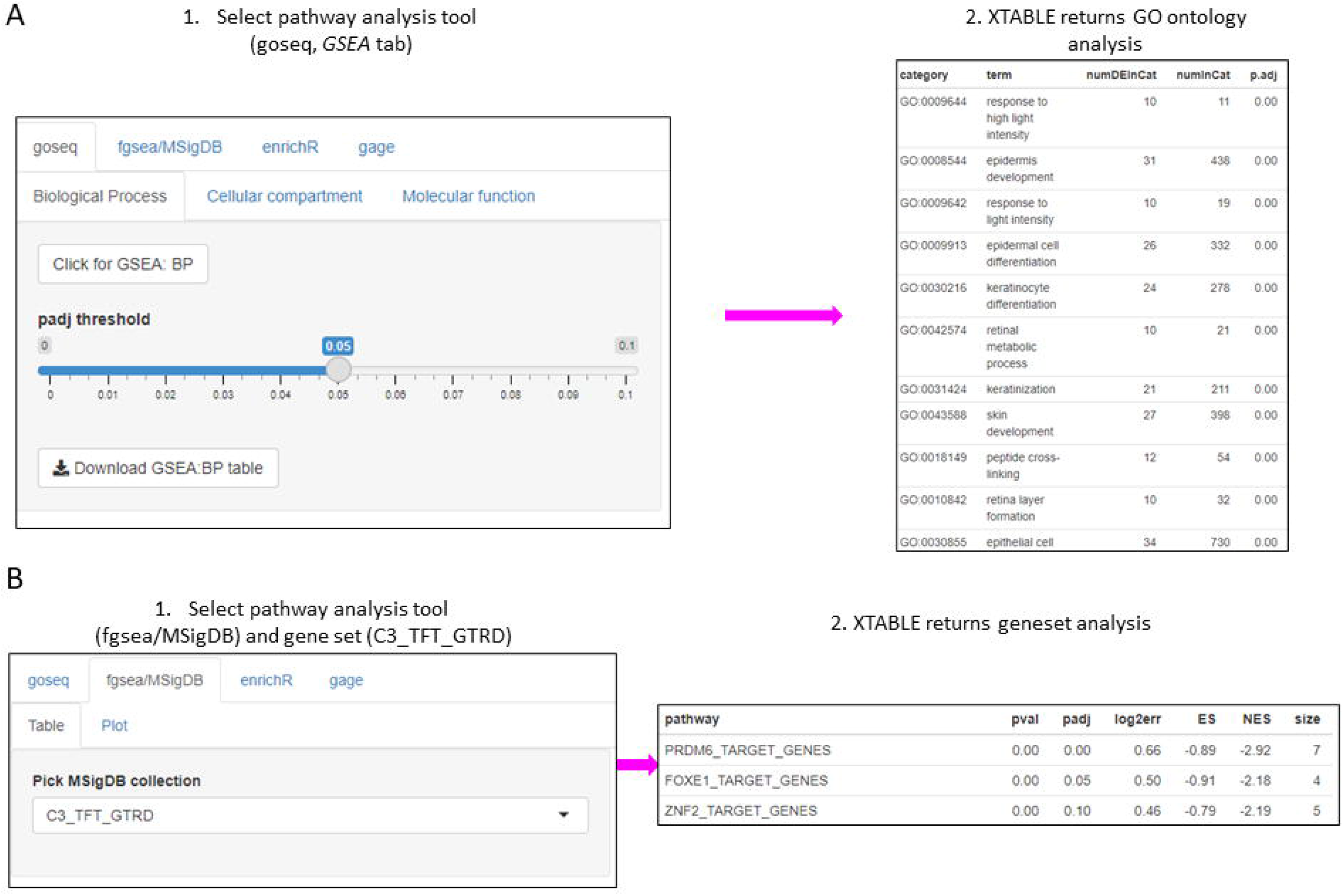
Gene-set enrichment analyses in a list differentially expressed genes using the *GSEA* tab. **A**. Gene-set enrichment analysis using the goseq tool of a list of differentially expressed genes obtained in the *DEG* tab. One of the three main GO ontologies can be selected for analysis at a time. After selection of a p-value, XTABLE returns a downloadable list of GO ontologies with associated statistics. **B**. Gene-set enrichment analysis using the fgsea/MSigDB tool. This tool allows the selection of any collection included in MSigDB and returns a list of signatures with associated statistics. The example shows the selection of the C3_TFT_GTRD collection (Transcription Factor Targets annotated in the Gene Transcription Regulation Database).

### Gene-centered analysis and user-defined transcriptional signatures

Users can investigate their own gene or group of genes of interest in the application. To facilitate this type of gene-centered analyses, we have included the *Genes* and *Signature* functions in XTABLE.

In the *Genes* tab, the user can analyse the expression of one gene of interest. This tab is divided in three tools. The ‘Expr’ tool returns a downloadable list of the maximum (if multiple probes map to the same gene symbol) normalized expression values for the gene of interest in all samples (Figure 5A). The ‘Indv_Contrast’ function returns the DEG analysis results including fold change and statistical significance values for a given gene in all groups of samples compared with a predetermined reference group (Figure 5B). Sample grouping and reference group depend on the study. The ‘Mult_Contrast’ function works similarly to the ‘Indv_Contrast’ function but allows merging of up to four groups of samples and the reference group for statistical analysis can be determined by the user. The example in Figure 5C shows the evolution of MYC expression in four groups of samples that represent normal, low grade PMLs, high-grade PMLs and invasive carcinomas (Figure 5C). The fold changes and p-values are the result with the comparison with the ‘normal’ group (normal normofluorescent, normal hypofluorescent and hyperplasia) as the reference group (Figure 5C).

**Figure 5.**
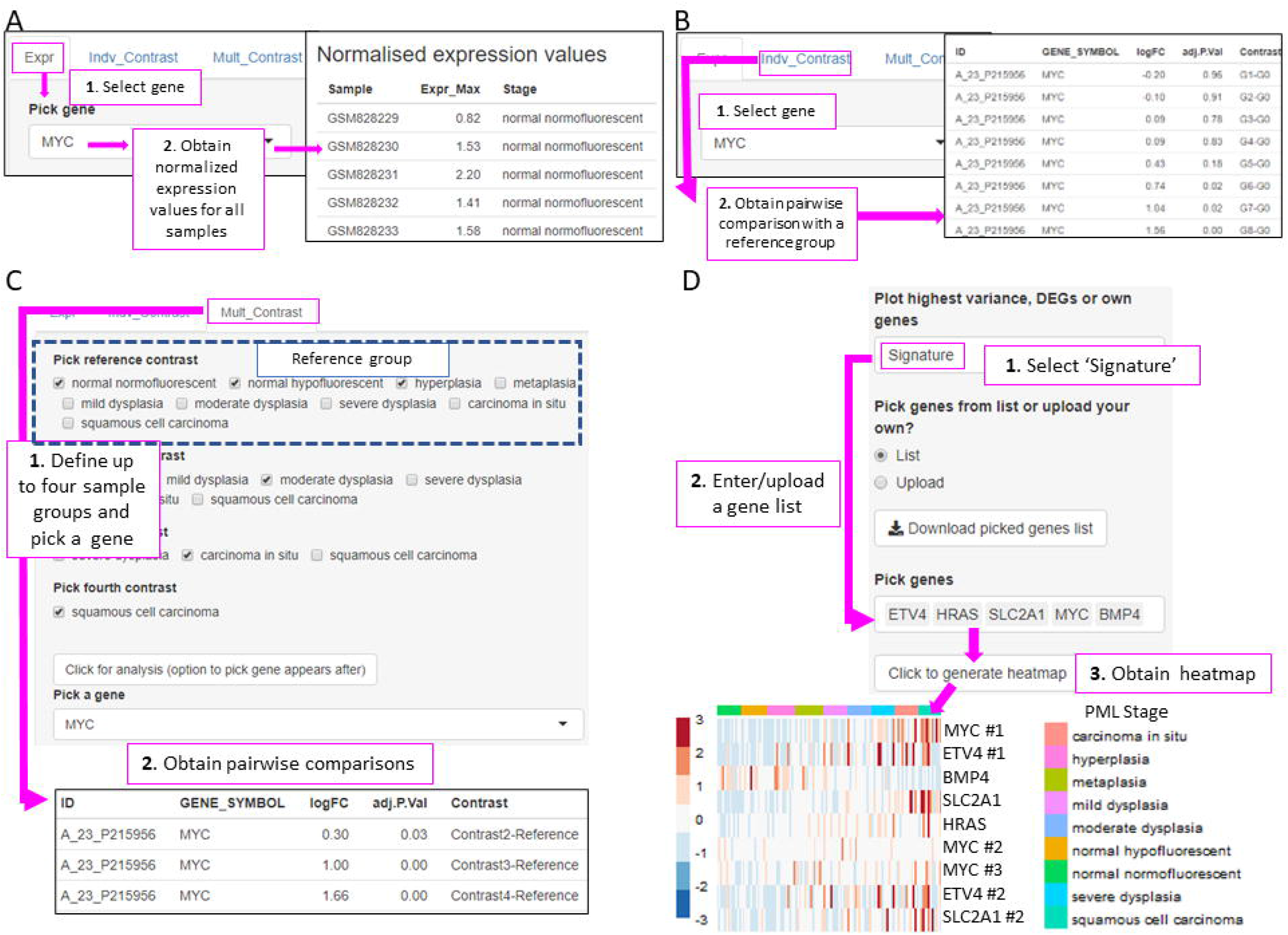
XTABLE functions to implement analyses on individual genes (*Gene* tab) and user defined gene signatures (*Signature* tab). **A**. The ‘Expr’ function (under the *Gene* tab) retrieves the normalized expression values for a gene of interest in all samples. **B**. The ‘Indiv_Contrast’ tool compares the expression of a gene of interest in groups of samples with a predetermined group. In the example, the function compares the expression of *MYC* in all stages with the normal normofluorescent group in GSE33479. **C**. The ‘Mult_Contrast’ tool enables the grouping of samples in up to four groups (contrasts) and statistical comparison with a reference group determined by the user. The example shows the analysis of *MYC* expression in four groups of samples from the GSE33479 cohort (normal, low-grade, high-grade, and invasive carcinomas). The ‘normal’ group is set as reference group for statistical analysis. **D**. Example of the use of the *Heatmap* tab to interrogate to visualize the expression of gene-sets in PMLs. Gene-sets can be defined by the user (as in the example) and are shown using the stage classification and entered manually or from a .csv file. Alternatively, the heatmap can be generated from a list of differentially expressed genes from the *DEG* tab or a selected number of genes filtered by variance. The three options can be selected in the scroll down menu. In the example shown, the heatmap shows all microarray probes associated to each gene symbol. P-values calculated using Welch’s t-test.

To facilitate the interrogation of biological processes driving PML progression, XTABLE also allows the interrogation of transcriptomic signatures using multiple functions. The *Signature* tab returns a list of normalized expression values and a graph with the signature scores (sum of log normalized expression values) for a gene-set determined by the user in all samples of the selected study.

The *Heatmap* tab returns a heatmap that displays the expression values in all samples in a selected study. The gene-set shown in the heatmap can be selected using three options from the scroll-down menu. With the ‘Signature’ option, the user can manually enter or upload a list of genes. The example in Figure 5D shows a heatmap generated from the GSE33479 dataset with 5 transcriptional targets of *SOX2*, an important LUSC driver. The ‘DEG’ option automatically selects the list of genes differentially expressed in the *DEG* tab. Finally, the ‘Variance’ option selects genes using a user-defined number of genes with the highest variance.

### Assessment of the correlation between chromosomal instability signatures and progression potential

Two studies have highlighted the potential role of CIN as predictor of low-grade (24) and high-grade (25) PMLs progression in LUSC. XTABLE can be used to explore this correlation in the four cohorts, identify genes and pathways altered by CIN and involved in driving it. Furthermore, XTABLE enables the user to carry out cross comparisons between cohorts to identify high-confidence signals. One interesting example of this cross comparisons is the correlation between CIN scores and progression in different cohorts using the *CIN-Score* and *ROC* tabs. Cohort GSE108124 focuses on CIS lesions with known progressive potential. As reported in that study (25), we found that the CIN5 signature segregates progressive and regressive lesions (AUC=1) (Figure 6A, Figure 6-figure supplement 1), whereas CIN70 and CIN25 signatures are somewhat poorer predictors of progression (Figure 6-figure supplement 2) (AUC= 0.82 and 0.81 respectively). Apart from the varied performance of different CIN-scores, this difference can also be attributed to the lack of data about some of the genes in the sequencing output. Furthermore, cohort GSE114489, which also contains dysplastic samples with known progression potential, demonstrates that although signatures are elevated in persistent dysplasias, neither CIN5 (AUC=0.72) (Figure 6B, Figure 6-figure supplement 3) nor CIN70 (AUC=0.74) and CIN25 (AUC=0.73) (Figure 6-figure-supplement 4) could accurately segregate persistent from regressive samples as efficiently as CIN5 in the GSE108124 cohort. Similarly, CIN scores did not accurately segregate regressive from persistent/progressive samples by progression status in the discovery cohort of GSE109743 (Figure 6C, Figure 6-figure supplement 5). A similar result was found with the validation cohort of GSE109743 (Figure 6-figure supplement 6). However, in cohorts with multiple stages represented (GSE109743 and GSE33479), there was a clear increase in CIN scores with increasing PML grade (Figures 6D and E, Figure 6-figure supplement 7), which is consistent with the increased risk of malignant progression in high-grade lesions. In summary, we found that CIN scores (specifically CIN5) are good predictors of malignant progression in CIS (GSE108124). In GSE114489, we observed an increase in the CIN5 score in persistent dysplasias but the discrimination between persistent and regressive dysplasias was poorer. In GSE109743, we did not observe significant differences between progressive/persistent PMLs and regressive lesions. This different performance of CIN scores in different cohorts can be attributed to multiple factors both technical and biological. The most important difference is that GSE108124 focuses on CIS lesions, a high-grade precursor of invasive LUSC, whereas the other cohorts focus on earlier lesions (GSE114489) or combinations of different stages (GSE109743). Additionally, microarray analysis was carried out with RNA extracted from microdissected PMLs in cohort GSE108124 providing an enriched epithelial signal.

**Figure 6:**
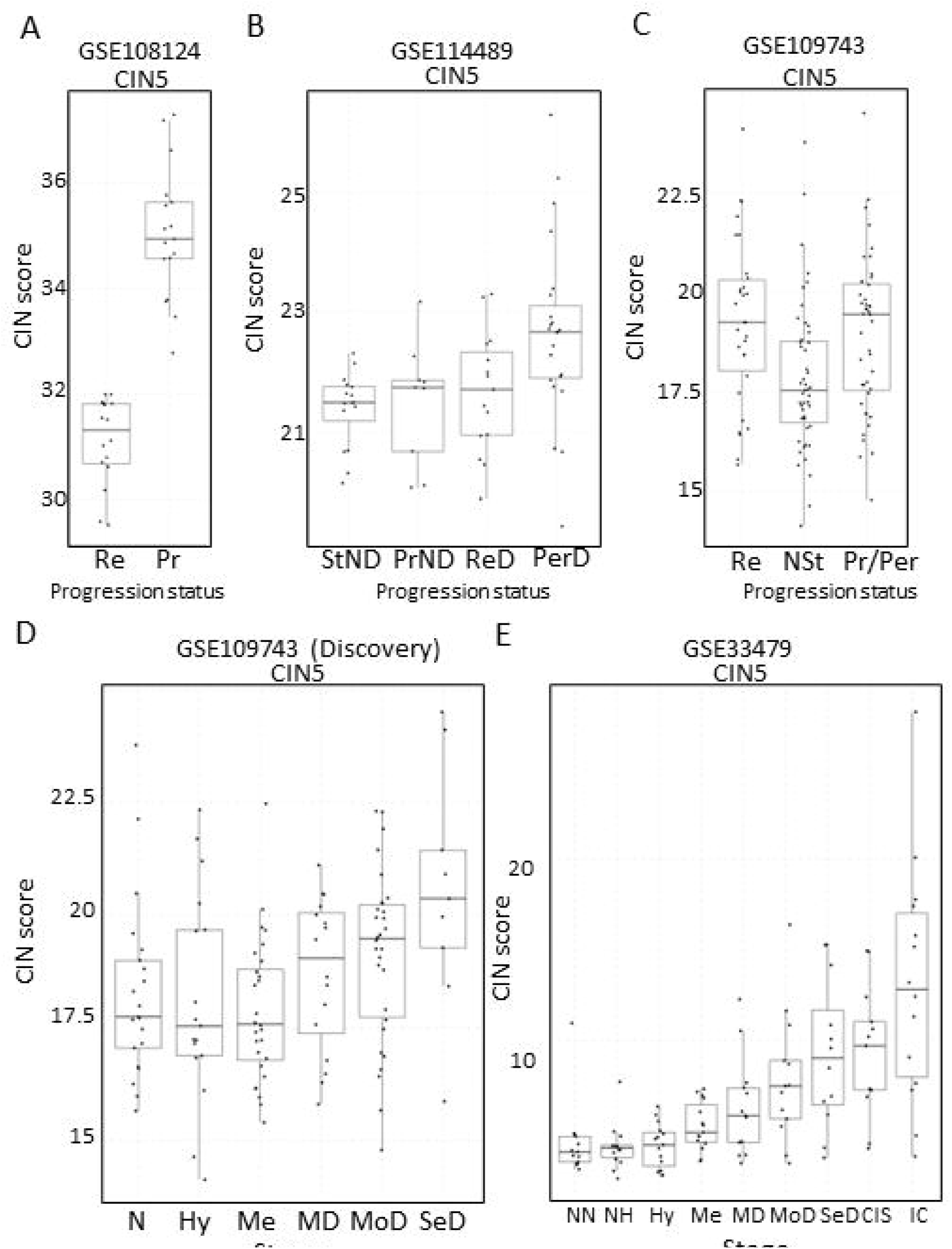
Association of CIN-scores with progression status and stage in the four cohorts of XTABLE. **A**. CIN5 score in regressive (Re) and progressive (Pr) CIS lesions from cohort GSE108124. **B**. CIN5 scores in stable non-dysplasias (StND), progressive non-dysplasias (PrND), regressive dysplasias (ReD) and persistent dysplasias (PerD) from cohort GSE114489. **C**. CIN5 scores in regressive (Re), normal stable (NSt) and progressive/persistent (Pr/Per) PMLs from cohort GSE109743). **D and E**. Evolution of CIN-scores in LUSC developmental stages for cohorts GSE109743 and GSE33479. N: normal; NN: normal normofluorescent; NH: normal hypofluorescent; Hy: Hyperplasia; Me:metaplasia; MD: mild dysplasia; MoD: moderate dysplasia; SeD: severe dysplasia; CIS: carcinoma in-situ; IC: invasive carcinoma. Boxplots show median and upper/lower quartile. Whiskers show the smallest and largest observations within 1.5xIQR.

### Mapping the evolution of the most relevant pathways involved in LUSC using XTABLE

Inactivation of the tumour suppressor genes *TP53* and *CDKN2A* (Figure 7A) are the most frequent somatic events in LUSC (6, 8). Other somatic alterations in driver genes are found at a lower frequency but often target the same pathways in different ways (Figure 7A) (6, 8). The squamous differentiation, PI3K/Akt and oxidative stress response are the most frequently targeted pathways in LUSC (8). The most frequent alteration targeting the squamous differentiation pathway is *SOX2* amplification, although inactivations of NOTCH proteins have also been proposed to target this pathway as they are mutually exclusive with *SOX2* amplification. A similar pattern of mutually exclusive somatic alterations targeting the same pathway has also been observed in the PI3K/Akt and oxidative stress response pathways (Figure 7A) (8). The role of these pathways in the transition between the LUSC developmental stages has not been addressed to date. The main reason for this is the paucity of genomic characterisations of PMLs and the lack of preclinical models of PMLs. Using XTABLE (*Signature* tab) to interrogate published transcriptional signatures correlated with these pathways can shed information about the stages at which they become active, and therefore what their potential role is in LUSC progression. To map changes in the activation of these three pathways to LUSC developmental stages, we use pre-designed transcriptional signatures from the MSigDB collections (32-34). Namely, the SOX2_BENPORATH (squamous differentiation) (35), HALLMARK_PI3K_AKT_MTOR_PATHWAY (PI3k/AKT pathway) and WP_NRF2_PATHWAY (oxidative stress response) signatures (Figure 1B, C and D respectively). When GSE33479 was interrogated, the signature scores of the three pathways increased significantly in either moderate or severe dysplasias when compared with normal normofluorescent mucosa. No significant increases were detected in mild dysplasias or earlier lesions. Similar results were observed with the GSE109743 cohort except for the WP_NRF2_PATHWAY (Figure 7-figure supplement 1). In this pathway, an increase of the signature was detected only in mild dysplasias compared with normal samples, and no significant changes were detected in any other PMLs stages.

**Figure 7:**
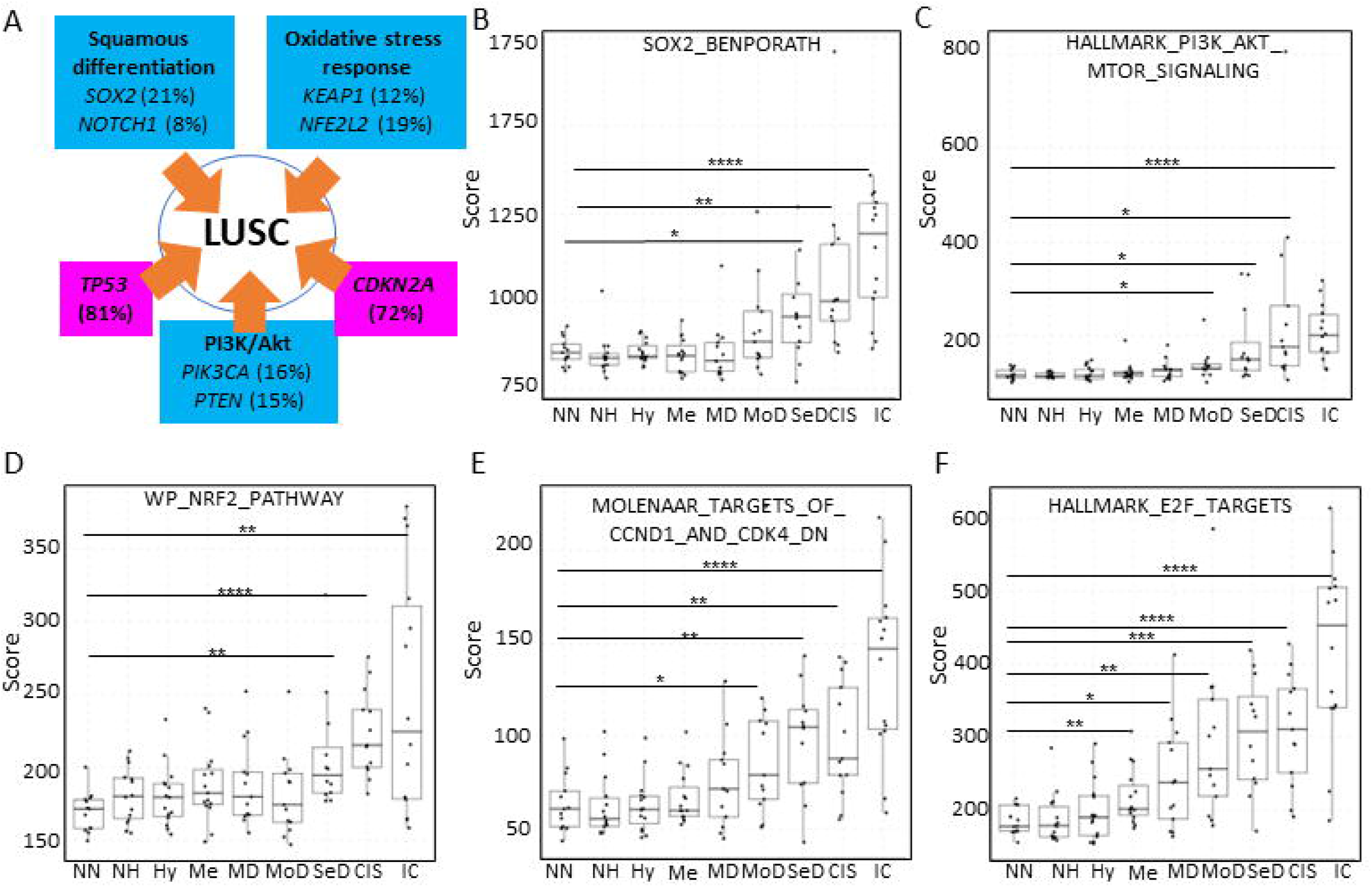
Mapping the evolution of the most relevant LUSC pathways to the LUSC developmental stages using published MSigDB transcriptional signatures. **A**. Diagram showing the most important pathways involved in LUSC and the genes involved in such pathways that are found genetically altered in LUSC tumours. **B**. Evolution of the SOX2 (the most frequent driver of the squamous differentiation pathway) transcriptional signature (SOX2_BENPORATH) during LUSC progression (GSE33479 cohort). **C**. Evolution of the PI3K/Akt pathway during LUSC progression (HALLMARK_PI3K_AKT_MTOR_SIGNALING). **D**. Evolution of the NRF2 (WP_NRF2_PATHWAY) transcriptional signature (correlated with the oxidative stress response) during LUSC progression. **E**. Evolution of a transcriptional signature correlated with cyclin-D1 and CDK4 (MOLENAAR_TARGETS_OF_CCND1_AND_CDK4_DN) during LUSC progression. *CDKN2A* alterations in LUSC lead to the inactivation of the p16INK^4a^, a CDK4 inhibitor. F: Evolution of the expression of E2F targets (HALLMARK_E2F_TARGETS). Inactivation of p16INK^4a^ results in activation of E2F transcriptional activity. Boxplots show median and upper/lower quartile. Whiskers show the smallest and largest observations within 1.5xIQR. *p<0.05, **p<0.01, ***p<0.001, p<0.0001 (Welch’s t-test).

We also interrogated the MSigDB collection to investigate the stages wherein the inactivation of *CDKN2A* is detected using an associated transcriptional signature. Since *CDKN2A* encodes the tumour suppressor p16INK^4a^, which inhibits the activity of the CDK4/Cyclin-D1 complexes and E2F transcriptional activity, activation of transcriptional signatures associated with these cell cycle regulators (MOLENAAR_TARGETS_OF_CCND1_AND_CDK4_DN and HALLMARK_E2F_TARGETS) can be used to monitor *CDKN2A* loss-of-function (36). The CDK4/Cyclin-D1 signature showed a significant increase in moderate dysplasias and later stages in cohort GSE33479 (Figure 7E) and in mild dysplasias and later stages in the GSE109743 (Figure 7-figure supplement 1). The increase of E2F signature was already detectable in metaplasias and later stages (Figure 7F) in the GSE33479 cohort, whereas in the GSE109743 cohort the increase in the E2F signature was observed starting in mild dysplasias (Figure 7-figure supplement 1).

Overall, these results show that the activity of squamous differentiation, PI3K/Akt and pathways start in the transition between high and low-grade lesions (Figure 1), typically moderate and severe dysplasias, indicating a role of these pathways in this transition, whereas their function in earlier stages (e.g., transition from normal epithelium to low-grade PMLs) is likely to be more limited. Our observations with the CDK4/Cyclin-D1 and E2F signatures indicate that the onset of this pathways start in earlier stages (metaplasias and mild-dysplasias).

## DISCUSSION

In this report, we have described XTABLE, an open source bioinformatic tool to explore gene expression in LUSC PMLs using four different transcriptomic datasets. The most novel aspect of this application is the emphasis on preinvasive disease (to our knowledge, the first bioinformatic tool focusing on PMLs), and the possibility of multiple sample stratifications by parameters that correlate with high-risk of malignant progression (stage, progression status and chromosomal instability). Treating lung cancer is complex. Chemotherapies, radiotherapy, targeted therapies, and immunotherapies save lives. However, the progress in lung cancer patient survival during the last 20 years is disappointing (16). Redirecting research efforts to prevent lung cancer and to detect its more treatable premalignant stages is the most efficient way to prevent lung cancer deaths to date. XTABLE offers researchers the possibility of accessing the most relevant transcriptomic databases on LUSC PMLs to assist in the understanding of PML biology and identify biomarkers for LUSC early detection. Biomarker identification, investigating the evolution of signaling pathways in multiple developmental LUSC stages, identification of immunomodulatory signals, changes in transcriptional signatures and exploring the causes and consequences of CIN in PMLs are amongst the multiple examples of promising applications of XTABLE for basic and translational biologists.

XTABLE also contributes to a more open, accessible, and inclusive science. Research laboratories often have restricted access to bioinformatic support due to funding constraints or lack of adequate collaborations. This limitation can be a major hurdle in the competitiveness of research groups as it may prevent hypothesis generation, validation of experimental results in patient cohorts or acquisition of preliminary results. XTABLE and similar applications can contribute to addressing those disadvantages with an accessible and versatile platform for gene-expression analysis. This application also contributes to the open science philosophy as it promotes the dissemination of data, accessibility, and transparency. Although XTABLE is unlikely to offer all the possibilities of analysis required by the scientific community, the application can be modified by the users to adapt it to their research questions. Additionally, XTABLE allows the download of results that can be subject to additional downstream analyses.

Our analysis of CIN-signatures and their correlation with the known progression potential of PMLs is one of the examples of the use of XTABLE to obtain a wider view of LUSC PML biology. CIN5 was a good predictor of CIS progression in the GSE108124 study, as already described by the authors of the study (25). However, CIN5 did not perform as well in other studies. Conceivably, the most plausible reason for this discrepancy is the nature of each study. GSE108124 focuses on CIS, the precursor lesions of invasive carcinomas and uses microdissected samples. The other two studies with progression status information focus on different stages and do not perform enrichment of the tumour component. GSE114489 investigates dysplasias, an earlier LUSC stage and lacks further subdivision into mild, moderate and severe dysplasic lesions. This limitation might result in data with more noise and poorer correlation with progression status. Nevertheless, this cohort did show an increase in CIN5 signature in persistent dysplasias when compared with regressive lesions, a difference that is not observed in GSE109743 (a cohort that does not separate PMLs by stage). These discrepancies point at several limitations of the use of CIN and its associated signatures as surrogates of progression potential. The use of CIN, and more specifically CIN-scores as a bona-fide predictor of progression might be limited to microdissected samples and CIS lesions. For example, the presence of tumour stroma could result in an underestimated CIN score. Additionally, CIS lesions are the most advanced premalignant stage and show the highest levels of CIN5 signatures, thereby contributing to a higher CIN-related signal. Comparisons with microdissected PMLs of earlier stages are necessary to address the extent of these limitations. Another limitation is the definition of progression and regression in different studies. Whereas GSE108124 provides a binary classification of PMLs (progressive and regressive), the other studies show a more complex classification that includes progressive and persistent lesions under the same category. Finally, the endpoint in the definition of the progression status also differs between studies. Specifically, the endpoint in GSE108124 is progression to invasive LUSC whereas the other studies define progression as transition to a higher grade. Conceivably, CIN might not play the same role in the transition between PMLs as it does in malignant transformation.

Regardless, CIN gene expression signatures present a very robust correlation with genomic instability indexes in cancer, and specifically in lung cancer (31). Although we have shown that the correlation of CIN signatures with progression potential varies between databases, they can be used to investigate other biological questions such as identification of CIN-tolerance and CIN-driver genes, identification of changes in the immune microenvironment associated with CIN (in light of the new role of CIN in modulating tumour immunity (37)) and interrogation of genes of interest for the user in CIN-high vs CIN-low PMLs.

Using XTABLE, we have mapped the most important signaling pathways targeted in LUSC to the developmental stages. We found that the onset activation of squamous differentiation, PI3K/Akt pathways occur in the transition from low to high-grade PMLs. Comprehensive genomic characterizations of PMLs have not been undertaken so far, and therefore, we are not able to map the onset of LUSC pathways with the genomic profiles of each premalignant stage. However, our observations suggest that the genomic alterations targeting those pathways should be more frequent in high-grade PMLs than low grade. Another explanation is that those mutations are present in low-grade PMLs but their effect on the pathways is only unleashed in the transition to high-grade lesions. Co-ocurring somatic alterations and microenvironment changes could explain this. Nevertheless, our results strongly indicate that squamous differentiation and PI3K/Akt pathways are unlikely to play a role in the earliest developmental stages (hyperplasias, metaplasias, and mild dysplasias). On the other hand, our observations with the CDK4/Cyclin-D1 and E2F signatures indicate an earlier onset (metaplasias and moderate dysplasias). Both signatures can be influenced by *CDKN2A*, a tumour suppressor gene involved in oncogene-induced senescence and inactivated in the vast majority of LUSC cases (38). It is conceivable that *CDKN2A* inactivation occur earlier than activation of oncogenic in order to prevent oncogene-induced senescence. For instance, *SOX2* overexpression has a negative effect in cell fitness in multiple experimental scenarios (39-42). Early inactivation of *CDKN2A* could be key to avoid this toxicity in LUSC. A similar scenario is also likely to occur with PI3K/Akt signaling, as this pathway needs to be exquisitely regulated to avoid senescence (43).

The activation of the squamous differentiation, PI3K/Akt pathways pathways in high-grade PMLs could be very useful as biomarkers to develop new modalities of early detection using multiple approaches. High-grade PMLs frequently undergo malignant progression (20) and therefore, detection of these lesions and removal is an optimal strategy to prevent deaths by LUSC. Design of appropriate molecular probes and theragnostic technologies that identify lesions with high levels pathway activation, and secreted proteins that are associated with those pathways are amongst the strategies to exploit pathway activation in early detection.

Finally, the aim of this article is not to prioritize any of the studies included in XTABLE, but to provide a tool for the simple and quick analysis of a large amount of biologically relevant data on PML biology. Each study has its own advantages and limitations. Therefore, the user is ultimately responsible for choosing the datasets that are more adequate to interrogate their research questions, compare results between databases considering the different scientific contexts of each study and interpret them in light of the different designs of each study.

## MATERIALS AND METHODS

### XTABLE download and installation

XTABLE can be downloaded from the GitLab repository (https://gitlab.com/cruk-mi/XTABLE) and it requires the previous installation of RStudio. Copy the commands, paste them on the RStudio console and run the command (Supplemental Video 1).

### XTABLE packages and construction

Bioconductor package GEOquery (2.54.1) was used to retrieve the data and the Bioconductor package Biobase (2.46.0) used to extract the gene expression values for microarray datasets. The Bioconductor package limma (3.42.2) was used to generate differentially expressed gene analysis results. Bioconductor packages AnnotationDbi (1.48.0) and org.Hs.eg.db (3.10.0) were used to retrieve additional gene IDs. Additional gene IDs were retrieved from Ensembl BioMart website (https://www.ensembl.org/biomart/martview/) with Ensembl Genes 104 dataset and Human Genes GRCh38.p13 to generate four mapping files with the following attributes: ‘Gene stable ID’ and ‘AGILENT WholeGenome 4×44k v1 probe’; ‘Gene stable ID’ and ‘Transcript stable ID’; ‘Gene stable ID’ and ‘Gene name’; ‘Gene stable ID’ and ‘NCBI gene (formerly Entrezgene) ID’ for GSE33479. For GSE114489, a file containing the attributes ‘Gene stable ID’ and ‘AFFY HuGene 1 0 st v1 probe’ was downloaded. The Bioconductor package edgeR (3.28.1) was used to calculate CPM values. R package pROC (1.17.0.1) was used to calculate AUC and generate ROC curves. Gene set enrichment analysis and pathway analysis was performed using Bioconductor packages ideal (1.10.0), fgsea (1.14) with MSigDB (https://www.gsea-msigdb.org/gsea/msigdb/collections.jsp) gene set collections (7.1), limma (3.42.2), pathview (1.26.0), enrichR (3.0), gage (2.36.0) with gageData (2.24.0), ReactomePA (1.30), progeny (1.8.0) and dorothea (0.99.0).

Deconvolution analysis was performed for microarray data using estimate (1.0.11) and for RNA-seq data using imsig (1.1.3). imsig requires filtering out genes with low variance which was performed with the package matrixStats (0.59.0). R package stats (3.6.0) used to generate principal component analysis data. ggplot2 (3.3.3), pheatmap (1.0.12), RColorBrewer (1.1-2) were used for making plots and ggpubr (0.4.0) was used to perform Welch’s t-test. tidyr (1.1.3), tibble (3.1.2), dplyr (2.0.6) and magrittr (2.0.1) were used for general data processing and formatting. Code was written using RStudio Workbench (1.4.1717.3) using R (4.0.3).

### Transcriptional signatures

To investigate the evolution of LUSC pathways, we downloaded the following gene sets from the Molecular Signatures Database (MSigDB v7.5.1):

BENPORATH_SOX2_TARGETS

(https://www.gsea-msigdb.org/gsea/msigdb/cards/BENPORATH_SOX2_TARGETS.html)

HALLMARK_PI3K_AKT_MTOR_SIGNALING (https://www.gsea-msigdb.org/gsea/msigdb/cards/HALLMARK_PI3K_AKT_MTOR_SIGNALING.html)

WP_NRF2_PATHWAY (https://www.gsea-msigdb.org/gsea/msigdb/cards/WP_NRF2_PATHWAY.html)

MOLENAAR_TARGETS_OF_CCND1_AND_CDK4_DN (https://www.gsea-msigdb.org/gsea/msigdb/cards/MOLENAAR_TARGETS_OF_CCND1_AND_CDK4_DN.html)

HALLMARK_E2F_TARGETS (https://www.gsea-msigdb.org/gsea/msigdb/cards/HALLMARK_E2F_TARGETS.html)

The signatures were manually curated to replace unrecognized gene symbols with alternative symbols used in each cohort. The signature scores returned by the XTABLE were calculated as the sum of log normalized expression values in each cohort.

## ACKNOWLEDGEMENTS

This work has been carried out with funding provided by the Cancer Research UK Lung Cancer Centre of Excellence (A25146) and the Cancer Research UK Manchester Institute (A27412) and the Manchester Biomedical Research Centre.

We would like to thank the Scientific Computing (SciCom) team of the CRUK Manchester Institute and Dr Anshuman Chaturvedi (The Christie Hospital) for their support in this project.

Representative images of premalignant stages in Figure 1 are shown with permission of the Manchester Cancer Research Centre (MCRC) Biobank, UK. The MCRC Biobank holds a generic ethics approval (Ref: 18/NW/0092) which can confer this approval to users of banked samples via the MCRC Biobank Access Policy.

## AUTHOR CONTRIBUTIONS

Conceptualization: MR, JO, JEB, CLG

Methodology: MR, MH, AK, JEB

Software: MR, MH, AK,

Validation: MR, JO, MH, AK, CD, JEB, CLG

Formal analysis: MR

Investigation: MR, JO, MH, AK, CD, JEB, CLG

Resources: MR, CLG

Data curation: MR, CLG

Writing (original draft preparation): MR, CLG

Writing (review and editing): MR, JO, AK, JEB, CLG

Supervision: AK, CD, CLG

Project administration: CLG

Funding acquisition: CD

## COMPETING INTERESTS

The authors do not have any competing interests to disclose.

## Notes

https://www.ncbi.nlm.nih.gov/geo/query/acc.cgi?acc=GSE33479

https://www.ncbi.nlm.nih.gov/geo/query/acc.cgi?acc=GSE109743

https://www.ncbi.nlm.nih.gov/geo/query/acc.cgi?acc=GSE114489

https://www.ncbi.nlm.nih.gov/geo/query/acc.cgi?acc=GSE108124

### Competing Interest Statement

The authors have declared no competing interest.

## REFERENCES

1. International Agency for Research on Cancer. Cancer Incidence in Five Continents Volume XI. 2021.

2. Torre LA, Siegel RL, Jemal A. Lung Cancer Statistics. Adv Exp Med Biol. 2016;893:1–19.

3. Hirsch FR, Suda K, Wiens J, Bunn PA, Jr. New and emerging targeted treatments in advanced non-small-cell lung cancer. Lancet. 2016;388(10048):1012–24.

4. Khuder SA. Effect of cigarette smoking on major histological types of lung cancer: a meta-analysis. Lung Cancer. 2001;31(2-3):139–48.

5. Cancer Genome Atlas Research N. Comprehensive molecular profiling of lung adenocarcinoma. Nature. 2014;511(7511):543–50.

6. Jamal-Hanjani M, Wilson GA, McGranahan N, Birkbak NJ, Watkins TBK, Veeriah S, et al. Tracking the Evolution of Non-Small-Cell Lung Cancer. N Engl J Med. 2017;376(22):2109–21.

7. Campbell JD, Alexandrov A, Kim J, Wala J, Berger AH, Pedamallu CS, et al. Distinct patterns of somatic genome alterations in lung adenocarcinomas and squamous cell carcinomas. Nat Genet. 2016;48(6):607–16.

8. Cancer Genome Atlas Research N. Comprehensive genomic characterization of squamous cell lung cancers. Nature. 2012;489(7417):519–25.

9. Kim Y, Hammerman PS, Kim J, Yoon JA, Lee Y, Sun JM, et al. Integrative and comparative genomic analysis of lung squamous cell carcinomas in East Asian patients. J Clin Oncol. 2014;32(2):121–8.

10. Mok TSK, Wu YL, Kudaba I, Kowalski DM, Cho BC, Turna HZ, et al. Pembrolizumab versus chemotherapy for previously untreated, PD-L1-expressing, locally advanced or metastatic non-small-cell lung cancer (KEYNOTE-042): a randomised, open-label, controlled, phase 3 trial. Lancet. 2019;393(10183):1819–30.

11. Paz-Ares L, Luft A, Vicente D, Tafreshi A, Gumus M, Mazieres J, et al. Pembrolizumab plus Chemotherapy for Squamous Non-Small-Cell Lung Cancer. N Engl J Med. 2018;379(21):2040–51.

12. Weinberg F, Gadgeel S. Combination pembrolizumab plus chemotherapy: a new standard of care for patients with advanced non-small-cell lung cancer. Lung Cancer (Auckl). 2019;10:47–56.

13. Targeted Drugs Fall Short in Squamous Lung Cancer. Cancer Discov. 2021;11(1):OF3.

14. Middleton G, Fletcher P, Popat S, Savage J, Summers Y, Greystoke A, et al. The National Lung Matrix Trial of personalized therapy in lung cancer. Nature. 2020;583(7818):807–12.

15. Redman MW, Papadimitrakopoulou VA, Minichiello K, Hirsch FR, Mack PC, Schwartz LH, et al. Biomarker-driven therapies for previously treated squamous non-small-cell lung cancer (Lung-MAP SWOG S1400): a biomarker-driven master protocol. Lancet Oncol. 2020;21(12):1589–601.

16. Surveillance, Epidemiology, and End Results (SEER) Program (http://www.seer.cancer.gov) SEER*Stat Database: Mortality - All COD, Aggregated With State, Total U.S. (1969-2018) <Katrina/Rita Population Adjustment>, National Cancer Institute, DCCPS, Surveillance Research Program, released May 2020. Underlying mortality data provided by NCHS (http://www.cdc.gov/nchs).

17. Crosbie PA, Balata H, Evison M, Atack M, Bayliss-Brideaux V, Colligan D, et al. Second round results from the Manchester ‘Lung Health Check’ community-based targeted lung cancer screening pilot. Thorax. 2019;74(7):700–4.

18. Crosbie PA, Balata H, Evison M, Atack M, Bayliss-Brideaux V, Colligan D, et al. Implementing lung cancer screening: baseline results from a community-based ‘Lung Health Check’ pilot in deprived areas of Manchester. Thorax. 2019;74(4):405–9.

19. National Lung Screening Trial Research T, Aberle DR, Adams AM, Berg CD, Black WC, Clapp JD, et al. Reduced lung-cancer mortality with low-dose computed tomographic screening. N Engl J Med. 2011;365(5):395–409.

20. Ishizumi T, McWilliams A, MacAulay C, Gazdar A, Lam S. Natural history of bronchial preinvasive lesions. Cancer Metastasis Rev. 2010;29(1):5–14.

21. Kadara H, Scheet P, Wistuba, II, Spira AE. Early Events in the Molecular Pathogenesis of Lung Cancer. Cancer Prev Res (Phila). 2016;9(7):518–27.

22. Thakrar RM, Pennycuick A, Borg E, Janes SM. Preinvasive disease of the airway. Cancer Treat Rev. 2017;58:77–90.

23. Pennycuick A, Teixeira VH, AbdulJabbar K, Raza SEA, Lund T, Akarca AU, et al. Immune Surveillance in Clinical Regression of Preinvasive Squamous Cell Lung Cancer. Cancer Discov. 2020;10(10):1489–99.

24. van Boerdonk RA, Daniels JM, Snijders PJ, Grunberg K, Thunnissen E, van de Wiel MA, et al. DNA copy number aberrations in endobronchial lesions: a validated predictor for cancer. Thorax. 2014;69(5):451–7.

25. Teixeira VH, Pipinikas CP, Pennycuick A, Lee-Six H, Chandrasekharan D, Beane J, et al. Deciphering the genomic, epigenomic, and transcriptomic landscapes of pre-invasive lung cancer lesions. Nat Med. 2019;25(3):517–25.

26. Merrick DT, Gao D, Miller YE, Keith RL, Baron AE, Feser W, et al. Persistence of Bronchial Dysplasia Is Associated with Development of Invasive Squamous Cell Carcinoma. Cancer Prev Res (Phila). 2016;9(1):96–104.

27. Beane JE, Mazzilli SA, Campbell JD, Duclos G, Krysan K, Moy C, et al. Molecular subtyping reveals immune alterations associated with progression of bronchial premalignant lesions. Nat Commun. 2019;10(1):1856.

28. Mascaux C, Angelova M, Vasaturo A, Beane J, Hijazi K, Anthoine G, et al. Immune evasion before tumour invasion in early lung squamous carcinogenesis. Nature. 2019;571(7766):570–5.

29. Guibert N, Mhanna L, Droneau S, Plat G, Didier A, Mazieres J, et al. Techniques of endoscopic airway tumor treatment. J Thorac Dis. 2016;8(11):3343–60.

30. Merrick DT, Edwards MG, Franklin WA, Sugita M, Keith RL, Miller YE, et al. Altered Cell-Cycle Control, Inflammation, and Adhesion in High-Risk Persistent Bronchial Dysplasia. Cancer Res. 2018;78(17):4971–83.

31. Carter SL, Eklund AC, Kohane IS, Harris LN, Szallasi Z. A signature of chromosomal instability inferred from gene expression profiles predicts clinical outcome in multiple human cancers. Nat Genet. 2006;38(9):1043–8.

32. Subramanian A, Tamayo P, Mootha VK, Mukherjee S, Ebert BL, Gillette MA, et al. Gene set enrichment analysis: a knowledge-based approach for interpreting genome-wide expression profiles. Proc Natl Acad Sci U S A. 2005;102(43):15545–50.

33. Liberzon A, Subramanian A, Pinchback R, Thorvaldsdottir H, Tamayo P, Mesirov JP. Molecular signatures database (MSigDB) 3.0. Bioinformatics. 2011;27(12):1739–40.

34. Liberzon A, Birger C, Thorvaldsdottir H, Ghandi M, Mesirov JP, Tamayo P. The Molecular Signatures Database (MSigDB) hallmark gene set collection. Cell Syst. 2015;1(6):417–25.

35. Ben-Porath I, Thomson MW, Carey VJ, Ge R, Bell GW, Regev A, et al. An embryonic stem cell-like gene expression signature in poorly differentiated aggressive human tumors. Nat Genet. 2008;40(5):499–507.

36. Molenaar JJ, Ebus ME, Koster J, van Sluis P, van Noesel CJ, Versteeg R, et al. Cyclin D1 and CDK4 activity contribute to the undifferentiated phenotype in neuroblastoma. Cancer Res. 2008;68(8):2599–609.

37. Tijhuis AE, Johnson SC, McClelland SE. The emerging links between chromosomal instability (CIN), metastasis, inflammation and tumour immunity. Mol Cytogenet. 2019;12:17.

38. Serrano M, Lin AW, McCurrach ME, Beach D, Lowe SW. Oncogenic ras provokes premature cell senescence associated with accumulation of p53 and p16INK4a. Cell. 1997;88(5):593–602.

39. Cho YY, Kim DJ, Lee HS, Jeong CH, Cho EJ, Kim MO, et al. Autophagy and cellular senescence mediated by Sox2 suppress malignancy of cancer cells. PLoS One. 2013;8(2):e57172.

40. Correia LL, Johnson JA, McErlean P, Bauer J, Farah H, Rassl DM, et al. SOX2 Drives Bronchial Dysplasia in a Novel Organotypic Model of Early Human Squamous Lung Cancer. Am J Respir Crit Care Med. 2017;195(11):1494–508.

41. Cox JL, Wilder PJ, Desler M, Rizzino A. Elevating SOX2 levels deleteriously affects the growth of medulloblastoma and glioblastoma cells. PLoS One. 2012;7(8):e44087.

42. Wuebben EL, Wilder PJ, Cox JL, Grunkemeyer JA, Caffrey T, Hollingsworth MA, et al. SOX2 functions as a molecular rheostat to control the growth, tumorigenicity and drug responses of pancreatic ductal adenocarcinoma cells. Oncotarget. 2016;7(23):34890–906.

43. Astle MV, Hannan KM, Ng PY, Lee RS, George AJ, Hsu AK, et al. AKT induces senescence in human cells via mTORC1 and p53 in the absence of DNA damage: implications for targeting mTOR during malignancy. Oncogene. 2012;31(15):1949–62.

