## Supplementary material for "Interrogating the Precancerous Evolution of Pathway Dysfunction in Lung Squamous Cell Carcinoma Using XTABLE": Figure supplement legends

**Figure 2-figure supplement 1:** Sample selection options for cohort GSE109743.

**Figure 2-figure supplement 2:** Visualization of CIN-scores in with XTABLE. The example shows CIN70 scores in GSE109743 cohort. The CIN signature can be changes in the ‘Pick signature’ option and the sample classification used in the x-axis can be selected with the “Plot x-axis” option.

**Figure 2-figure supplement 3:** Example of ROC curves visualization for a gene of interest (NRTK2) in PML samples stratified by low and high-grade.

**Figure 4-figure supplement 1:** Example of pathway analysis (*PA* tab) output for a gene list obtained with the *DEG* function.

**Figure 6-figure supplement 1:** ROC analysis of CIN5 as predictor of CIS progression in the GSE108124 cohort.

**Figure 6-figure supplement 2:** Analysis of CIN70 and CIN25 scores as predictors of CIS progression in GSE108124.

**Figure 6-figure supplement 3:** ROC analysis of CIN5 as predictor of PML progression in the GSE114489 cohort.

**Figure 6-figure supplement 4:** Analysis of CIN70 and CIN25 scores as predictors of PML progression in GSE114489.

**Figure 6-figure supplement 5:** Analysis of CIN70, CIN25 and CIN5 scores as predictors of PML progression in GSE109743.

**Figure 6-figure supplement 6:** CIN5 scores in the GSE109743 cohort with samples classified by progression status.

**Figure 6-figure supplement 7:** Evolution of CIN5 scores by PML stage in the validation cohort of GSE109743.

**Figure 7-figure supplement 1:** Evolution of five transcriptional signatures in cohort GSE109743.
