## Supplementary figures and images for "Interrogating the Precancerous Evolution of Pathway Dysfunction in Lung Squamous Cell Carcinoma Using XTABLE"

### Figure 2-figure supplement 1

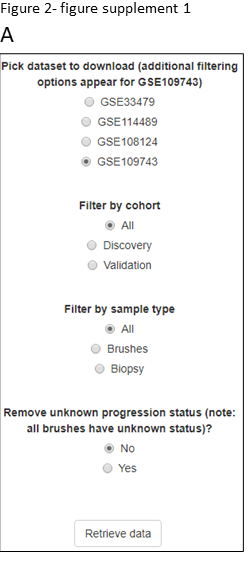

### Figure 2-figure supplement 2

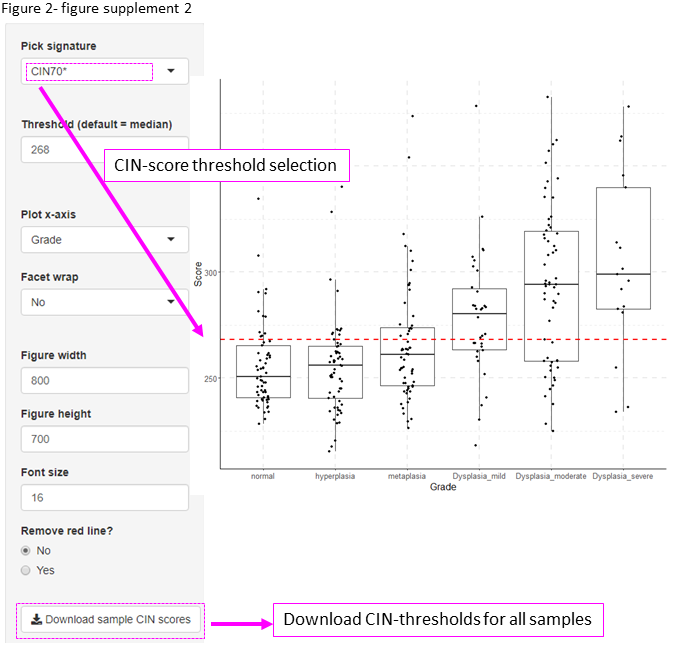

### Figure 2-figure supplement 3

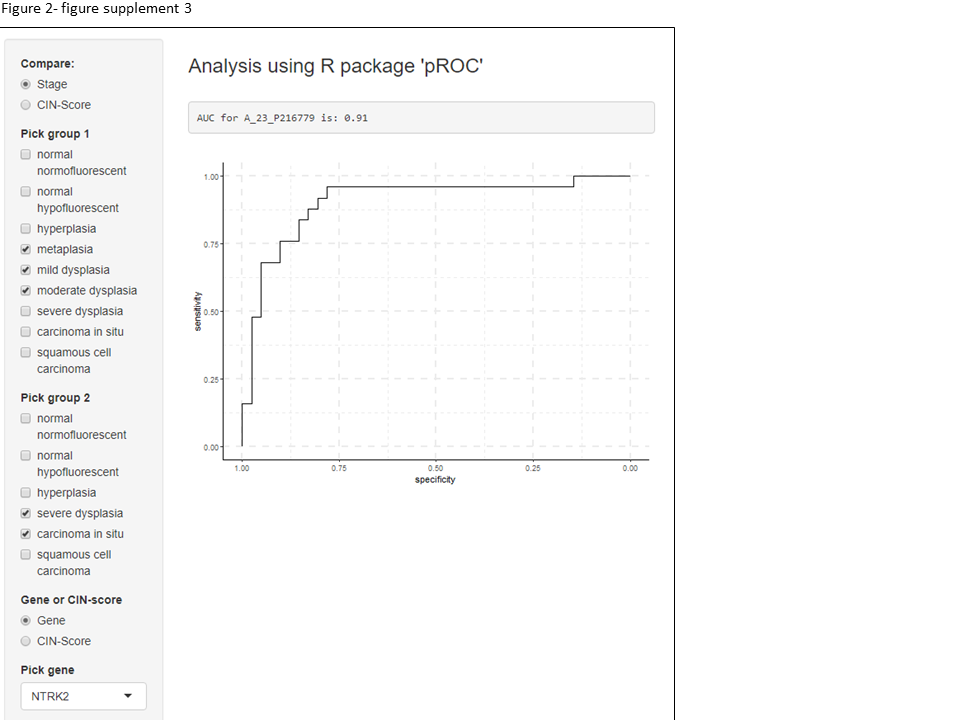

### Figure 4-figure supplement 1

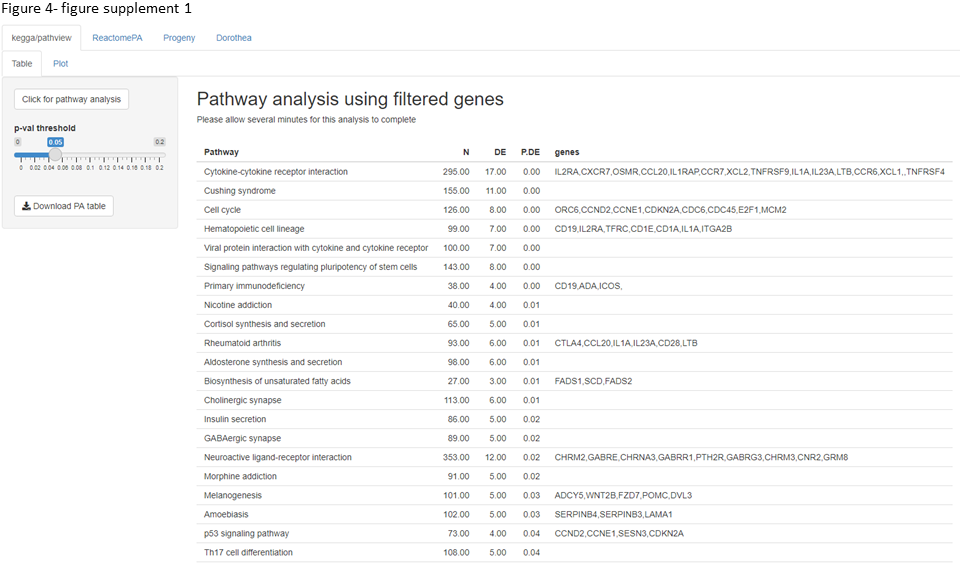

### Figure 6-figure supplement 1

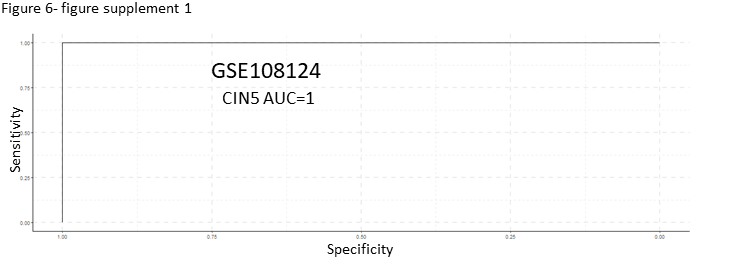

### Figure 6-figure supplement 2

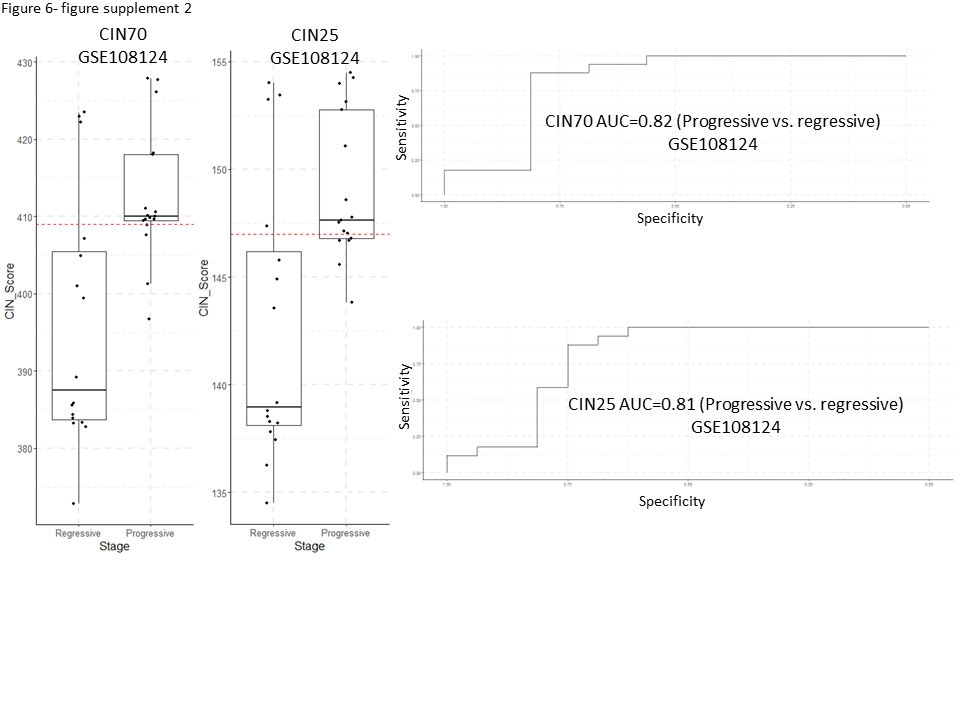

### Figure 6-figure supplement 3

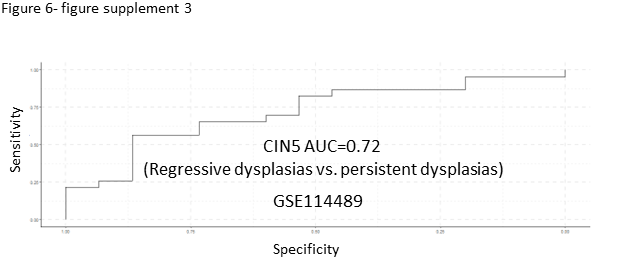

### Figure 6-figure supplement 4

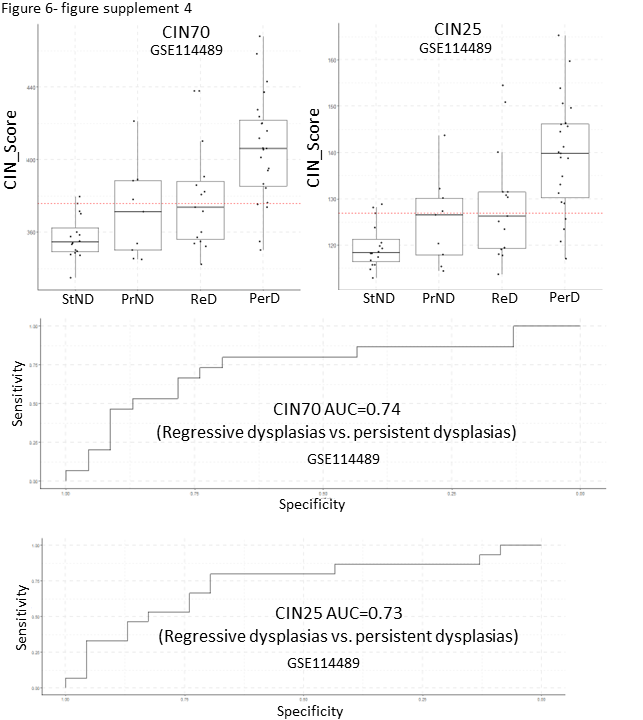

### Figure 6-figure supplement 5

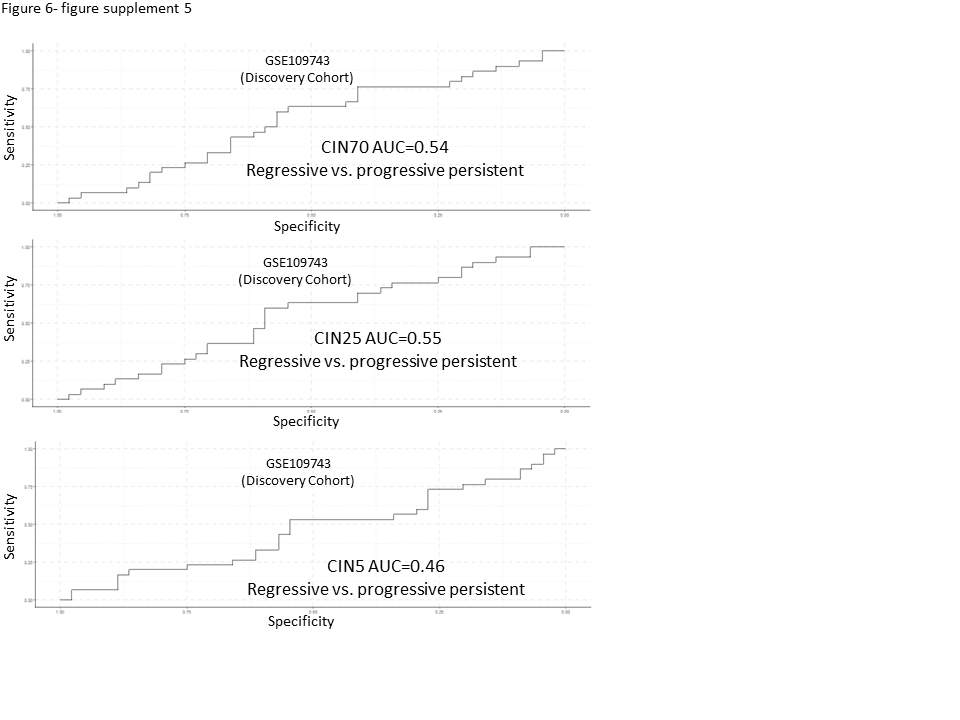

### Figure 6-figure supplement 6

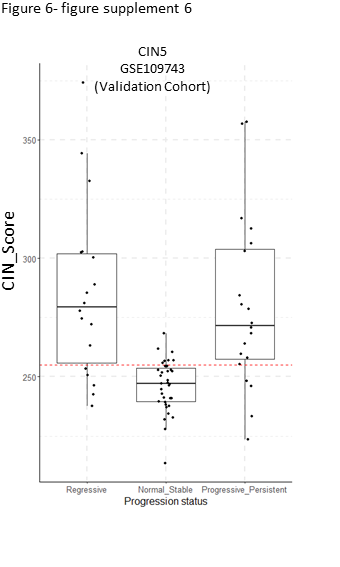

### Figure 6-figure supplement 7

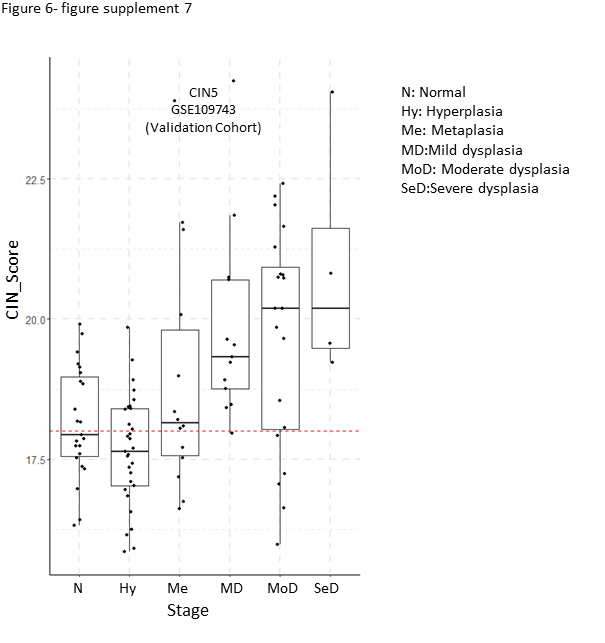

### Figure 7-figure supplement 1

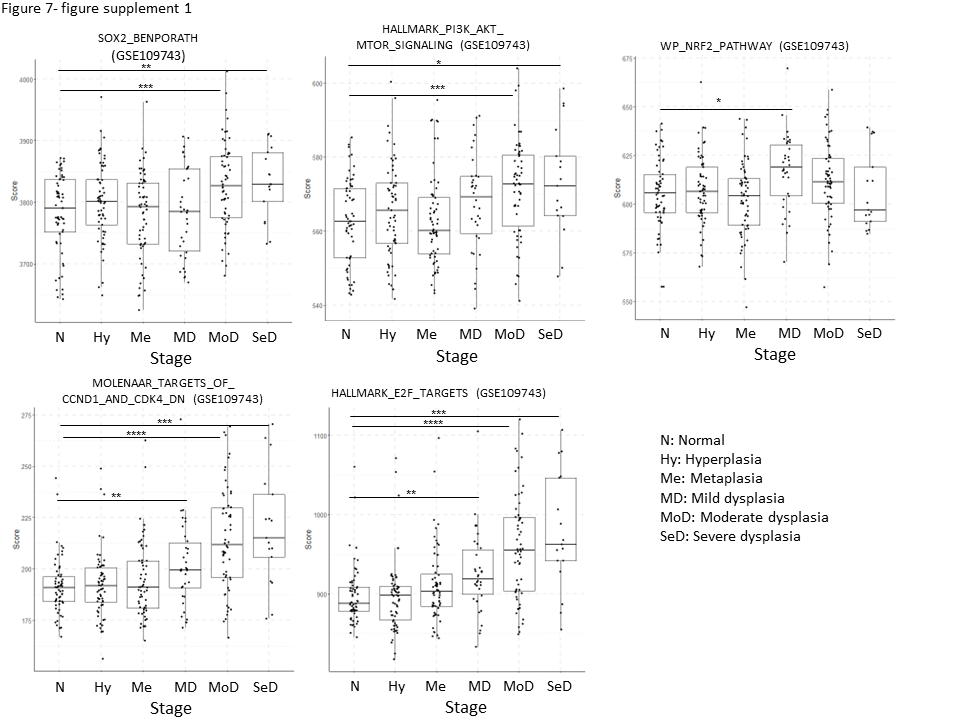
